# Neural responses to repeated noise structure in sounds are invariant to temporal interruptions

**DOI:** 10.1101/2023.02.22.529572

**Authors:** Björn Herrmann

**Affiliations:** Rotman Research Institute, Baycrest, M6A 2E1, North York, ON, Canada, Department of Psychology, University of Toronto, M5S 1A1, Toronto, ON, Canada

**Keywords:** Electroencephalography, frozen noise, temporal regularity, auditory perception, neural synchronization

## Abstract

The ability to extract meaning from acoustic environments requires sensitivity to repeating sound structure. Yet, how events that repeat are encoded and maintained in the brain and how the brain responds to events that reoccur at later points in time is not well understood. In two electroencephalography experiments, participants listened to a longer, ongoing white-noise sound which comprised shorter, frozen noise snippets that repeated at a regular 2-Hz rate. In several conditions, the snippet repetition discontinued for a brief period after which the noise snippet reoccurred. The experiments aimed to answer whether neural activity becomes entrained by the regular repetition of noise snippets, whether entrained neural activity self-sustains during the discontinuation period, and how the brain responds to a reoccurring noise snippet. Results show that neural activity is entrained by the snippet repetition, but there was no evidence for self-sustained neural activity during the discontinuation period. However, auditory cortex responded with similar magnitude to a noise snippet reoccurring after a brief discontinuation as it responded to a noise snippet for which the snippet repetition had not been discontinued. This response invariance was observed for different onset times of the reoccurring noise snippet relative to the previously established regularity. The results thus demonstrate that the auditory cortex sensitively responds to, and thus maintains a memory trace of, previously learned acoustic noise independent of temporal interruptions.

## Introduction

The sounds humans encounter in everyday life, such as speech and music, are rich of structured amplitude and frequency components that repeat over time, and humans are sensitive to such repetitions (McDermott et al., 2013; Teki et al., 2013; Sohoglu and Chait, 2016; Herrmann et al., 2020). Sensitivity to the repetition of sound structure is thought to support generating predictions about future sounds (Jones and Boltz, 1989; Barnes and Jones, 2000; Bendixen, 2014; Henry and Herrmann, 2014; Nobre and van Ede, 2018), enabling acoustic-change detection (Schröger, 2007; Winkler et al., 2009), and facilitating the perception of speech (Idemaru and Holt, 2011; Giraud and Poeppel, 2012; Peelle and Davis, 2012; Baese-Berk et al., 2014). Yet, how events that repeat are encoded and maintained in the brain and how the brain responds to events when they reoccur is not fully understood.

A variety of different approaches have been used to investigate the perception and neural correlates of repeating structure in sounds (Guttman and Julesz, 1963; Warren et al., 2001; Schröger, 2005; Agus et al., 2010; Agus and Pressnitzer, 2013; Bendixen, 2014; Andrillon et al., 2015; Barascud et al., 2016; Teki et al., 2016; Viswanathan et al., 2016; Andrillon et al., 2017; Kang et al., 2017; Bianco et al., 2020; Herrmann et al., 2021; Hodapp and Grimm, 2021; Dauer et al., 2022; Ringer et al., 2022). One fruitful line of research has focused on acoustic noise as stimuli to investigate how individuals perceive and learn repeating structure in sounds (Warren et al., 2001; Kaernbach, 2004; Agus et al., 2010; Agus and Pressnitzer, 2013; Luo et al., 2013; Andrillon et al., 2015; Rajendran et al., 2016; Andrillon et al., 2017; Dauer et al., 2022). The use of acoustic noise to examine the sensitivity to repeated structure has the advantage that a listener will not have previously heard the specific instantiation of a noise sound and will not be able to use semantic category labels because each noise consists of randomly generated numbers. Although the specific paradigms and stimuli have varied in previous studies, in a subset of studies, a repeated, short snippet of noise (∼0.2 s) was embedded in a longer, ongoing noise stimulus (Andrillon et al., 2015; Andrillon et al., 2017; Dauer et al., 2022). Both the short noise snippet and the longer noise are generated based on the same random-number distribution (e.g., Gaussian white noise). The short noise snippet is sometimes referred to as “frozen noise” because the exact array of random values that makes up the snippet are “frozen” in place and repeated within the ongoing noise stimulus (Warren et al., 2001; Andrillon et al., 2015; Rajendran et al., 2016; Dauer et al., 2022). The ongoing noise, in contrast, consists of nonrepeating random values drawn from the same distribution (Figure 1). The repetition of the short noise snippet in the ongoing noise forms a regular auditory pattern that a listener can perceive.

**Figure 1:**
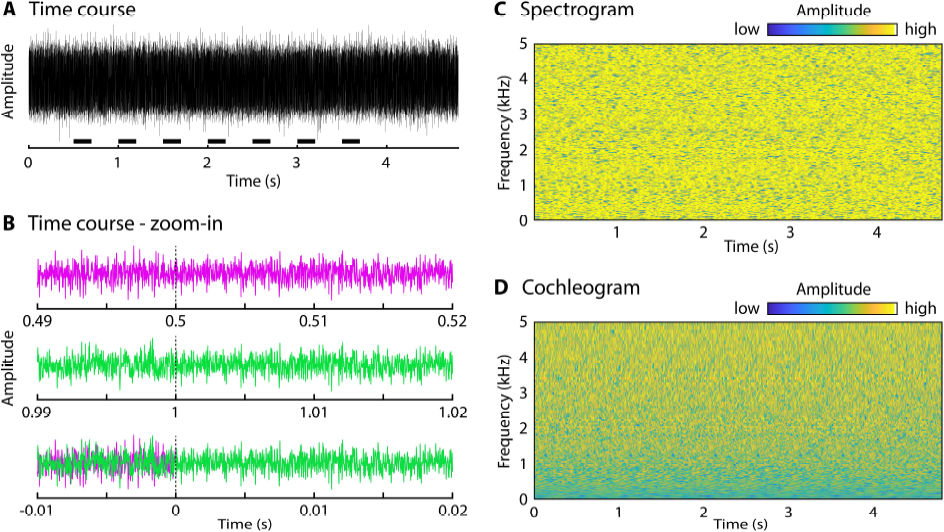
Sample noise stimulus. A: Time course of a sample noise stimulus with seven embedded noise snippets. The seven short black lines above the x-axis indicate the onset and the 0.2-s duration of each noise snippet. B: 30-ms zoomed-in time courses time-locked to the onset of the first (onset at 0.5 s; top panel) and second (onset at 1 s; middle panel) embedded noise snippet. The bottom panel shows the same time courses of the snippet again, overlaid and time-locked to snippet onset. The overlaid time courses demonstrate that snippets are identical (purple line and green line overlap), but that the noise stimulus differs prior to snippet onset. C: Spectrogram of the sample stimulus in panel A. D: Cochleogram of sample stimulus in panel A (cochleogram was calculated using procedures described in previous work: McDermott and Simoncelli, 2011). The time course (A), spectrogram (C), and cochleogram (D) show that no silences or other acoustic cues separate embedded snippet repetitions from the ongoing noise.

Individuals have remarkable memory for such frozen, repeated noise snippets that form a regular auditory pattern in an ongoing noise. Listeners are more likely to detect the presence of a pattern when the same, compared to a new, noise snippet is used on different trials (Andrillon et al., 2015; Andrillon et al., 2017; Dauer et al., 2022). Individuals further appear to have memory for specific auditory patterns even days and weeks after they have been exposed to them (Agus et al., 2010; Viswanathan et al., 2016; Bianco et al., 2020; Bianco et al., 2023). However, how the brain maintains memory representations for a noise snippet after it has been repeated is unclear. The current work focuses on short, within-second timescales to investigate the neural responses to a noise snippet reoccurring after a brief discontinuation of a snippet repetition.

Low-frequency neural activity is thought to reflect fluctuations in neuronal excitability (Bishop, 1933; Adrian and Matthews, 1934; Lakatos et al., 2005) that entrains to periodic repetitions of sounds or sound features (Picton et al., 2003; Stefanics et al., 2010; Herrmann et al., 2013; Herrmann et al., 2023) such that high excitability periods align with occurrences of the sounds or sound features to facilitate their processing (Thut et al., 2011; Henry and Herrmann, 2014; Lakatos et al., 2019; Obleser and Kayser, 2019). The term entrainment is used broadly here because responses to periodic repetitions of sounds could either reflect neural oscillations or evoked responses, and distinguishing between the two response types is empirically challenging (Klimesch et al., 2007; Sauseng et al., 2007; Capilla et al., 2011; Fellinger et al., 2011; Notbohm et al., 2016; Keitel et al., 2021).

Entrainment of neural activity has been observed for noise snippets that are repeated in an ongoing noise (Andrillon et al., 2015; Andrillon et al., 2017; Ringer et al., 2023b, a), providing a potential mechanism that enables the encoding of the noise snippet. Neural activity that is entrained by temporally regular stimulation – if oscillatory in nature – is hypothesized to self-sustain at the repetition rate when the stimulation discontinues (Pikovsky et al., 2001; Lakatos et al., 2013b; Henry and Herrmann, 2014; Keitel et al., 2022). Self-sustainability of entrained neural activity may facilitate the neural representation of a snippet even when its repetition temporarily discontinues and may support neural responses to a subsequent reoccurrence of the noise snippet. However, whether neural activity self-sustains after a snippet repetition discontinues and possibly facilitates responses to a reoccurring noise snippet is unclear. Moreover, if the entrained response reflects an evoked response, no self-sustainability of neural activity would be expected. Whether or not neural activity self-sustains may thus speak to the underlying response mechanism.

Perception of sounds and neural entrainment can depend on the timing at which events occur (Large and Jones, 1999; Barnes and Jones, 2000; Jones et al., 2002; Jones et al., 2006; Lange, 2010; Kayser et al., 2015; Herrmann et al., 2016; Morillon et al., 2016; Herrmann et al., 2023). For example, listeners detect auditory patterns made of repeating noise snippets in an ongoing noise better when the noise snippet is repeated at regular compared to irregular intervals (Rajendran et al., 2016; Dauer et al., 2022). Making perceptual inferences about a sound is also better when the sound occurs in-sync with a temporally regular stimulation – that is, at a time predicted by the periodicity of a regular stimulus repetition – compared to out-of-sync (Barnes and Jones, 2000; Jones et al., 2002; McAuley and Jones, 2003; Jones et al., 2006; Herrmann et al., 2016). The perceptual benefit of presenting a sound in-sync with a previous stimulus regularity is thought to arise from entrained, self-sustained neural activity (Large and Jones, 1999; Lakatos et al., 2013b; Henry and Herrmann, 2014), such that high-excitability periods occur at in-sync times and low-excitability periods at out-of-sync times. Perceptual sensitivity and neural responses to a reoccurring noise snippet may thus be facilitated if the snippet reoccurs in-sync compared to out-of-sync with a previously established temporal regularity, but this has not been investigated.

In two electroencephalography (EEG) experiments, the current study investigates the neural responses to frozen noise snippets that repeat at a regular rate. The experiments aim to answer whether neural activity self-sustains during a discontinuation of the repetition and how the brain responds to a reoccurring noise snippet. Specifically, Experiment 1 is concerned with the neural activity during the discontinuation period and the neural response to a reoccurring noise snippet. Experiment 2 is concerned with the degree of synchrony of a reoccurring noise snippet relative to a previously established stimulus regularity. The two EEG experiments will help understand how the brain maintains neural representations of recently learned structure in sounds.

## General methods

### Participants

Thirty-seven younger adults (18–34 years) participated in two EEG experiments of the current study. Each person participated only in one of the two experiments. Participants gave written informed consent prior to the experiment and were paid $7.5 CAD per half-hour for their participation. Participants reported having normal hearing abilities. The study was conducted in accordance with the Declaration of Helsinki, the Canadian Tri-Council Policy Statement on Ethical Conduct for Research Involving Humans (TCPS2-2014), and was approved by the Research Ethics Board of the Rotman Research Institute at Baycrest.

### Sound environment and stimulus presentation

Data collection was carried out in a sound-attenuating booth. Sounds were presented via Sennheiser (HD 25-SP II) headphones and a RME Fireface 400 external sound card. Stimulation was run using Psychtoolbox in MATLAB (v3.0.14; MathWorks Inc.) on a Lenovo T480 laptop with Microsoft Windows 7. Visual stimulation was projected into the sound booth via screen mirroring. All sounds were presented at a comfortable listening level that was fixed across participants (∼70-75 dB SPL).

### Acoustic stimulation and procedure

Stimulus generation was based on procedures described in previous work (Agus et al., 2010; Agus and Pressnitzer, 2013; Andrillon et al., 2015; Andrillon et al., 2017; Dauer et al., 2022). Each sound was a Gaussian white noise with a duration of either 4.8 s (Experiment 1) or 5 s (Experiment 2) sampled at 44.1 kHz. Participants listened either to white-noise sounds without additional characteristics – that is, they did not contain a pattern – or to white-noise sounds that had embedded a short, repeating white-noise snippet that created a regular pattern. In detail, a short noise snippet (mean duration across experiments: 0.253 s; range of durations: 0.079–0.411 s), generated from the same Gaussian white-noise distribution, was embedded in the 4.8-s or 5-s longer noise at six or more instances at a constant onset-to-onset interval of 0.5 s (2 Hz). A unique, newly created noise snippet was generated for each trial. The onset of the first embedded noise snippet was 0.5 s after the onset of the longer white noise. The specific number of repetitions depended on the experiment and condition, and is described for each experiment individually below. There were no silences or other acoustic cues separating embedded snippet repetitions from the ongoing noise, because snippets and ongoing noise were generated from the same Gaussian white-noise (i.e., random-number) distribution (Agus et al., 2010; Agus and Pressnitzer, 2013; Andrillon et al., 2015; Andrillon et al., 2017; Dauer et al., 2022). That no acoustic cues are present in the noise stimuli is also depicted in the time course, spectrogram, and cochleogram (McDermott and Simoncelli, 2011) of a sample noise stimulus that included a snippet repetition shown in Figure 1. Because of the absence of acoustic cues separating embedded snippet repetitions from the ongoing noise, perception of a pattern in these sounds required the detection of the noise-snippet repetition within the ongoing noise stimulus (Figure 1B).

During the experimental procedures, participants listened to the 4.8-s or 5-s white-noise sounds, one at the time, and were asked to judge whether a pattern – that is a noise-snippet repetition – was present or not. Participants were required to withhold their response until 0.1 s after sound offset where a happy and a sad emoticon smiley were presented visually side by side (Herrmann et al., 2011a; Herrmann et al., 2011b; Herrmann et al., 2021). If participants heard a pattern, they had to press the button for the happy smiley. If they did not hear a pattern, they had to press the button for the sad smiley. The position of the happy and sad smileys (left vs. right) was random across trials and participants did therefore not know which button to press during listening, that is, prior to the visual presentation (Herrmann et al., 2011a; Herrmann et al., 2011b; Herrmann et al., 2021). The next trial started 1.9 s after they had made a response.

Detection of a noise-snippet repetition depends on the duration of the noise snippet (Rajendran et al., 2016). In order to ensure that pattern detection was difficult but manageable, each participant underwent a threshold-estimation procedure at the beginning of the experimental session during which the duration of the noise snippet was estimated that would yield a detection rate for snippet repetitions of 0.75 (with a few exceptions detailed for each experiment below). To this end, participants listened to stimuli with embedded noise snippets varying at eight, equally-spaced durations: 0, 0.057, 0.114, 0.171, 0.229, 0.286, 0.343, 0.4 s (a duration of 0 means that no snippet repetition was embedded). Each snippet duration was presented 8 times in random order. This block lasted about 8 min. For each duration level, the proportion of ‘pattern present’ responses was calculated. A logistic function was fit to the proportion data and the duration corresponding to a proportion of 0.75 (with a few exceptions, see below) was calculated and used for the participant during the main experimental procedures.

### Analysis of behavioral data during main experimental procedures

A response was considered a hit if a participant responded ‘pattern present’ to a sound that contained a repetition of a noise snippet. A response was considered a false alarm if a person responded ‘pattern present’ to a sound that did not contain a repetition of a noise snippet. The hit rate was calculated for each of the experimental conditions (see below for each experiment), whereas one false alarm rate was calculated per participant (i.e., non-condition-specific). Because pattern-absent trials were presented among the conditions of pattern-present trials, no condition-specific false alarm rate could be calculated (cf. Agus et al., 2010; Agus and Pressnitzer, 2013; Andrillon et al., 2015; Andrillon et al., 2017; Kang et al., 2017; Hodapp and Grimm, 2021; Kang et al., 2021; Dauer et al., 2022; Ringer et al., 2022). False alarm and hit rates were used to calculate perceptual sensitivity (d-prime) for each participant and condition (Macmillan and Creelman, 2004) using the formula d-prime = z(hit rate) - z(false alarm rate), where z is the inverse of the normal cumulative distribution function (norminv in Matlab).

### Electroencephalography recordings and preprocessing

Electroencephalographical signals were recorded from 16 scalp electrodes (Ag/Ag–Cl-electrodes; 10-20 placement) and the left and right mastoids using a BioSemi system (Amsterdam, The Netherlands). The sampling frequency was 1024 Hz with an online low-pass filter of 208 Hz. Electrodes were referenced online to a monopolar reference feedback loop connecting a driven passive sensor and a common-mode-sense (CMS) active sensor, both located posteriorly on the scalp.

Offline analysis was conducted using MATLAB software. An elliptic filter was used to suppress power at the 60-Hz line frequency. Data were re-referenced by averaging the signal from the left and right mastoids and subtracting the average separately from each of the 16 channels. Data were filtered with a 0.7-Hz high-pass filter (length: 2449 samples, Hann window). The high-pass filter was specifically designed to provide strong DC suppression that avoids requiring baseline correction and removes slow drifts (Maess et al., 2006; Herrmann et al., 2011a; Herrmann et al., 2011b). Data were further low-pass filtered at 22 Hz (length: 211 samples, Kaiser window), divided into epochs ranging from 1 s prior to sound onset to 1 s post sound offset (−1 to 5.8 s for Experiment 1, and −1 to 6 s for Experiment 2; time-locked to sound onset), and downsampled to 512 Hz. Independent components analysis (runica method, Makeig et al., 1996; logistic infomax algorithm, Bell and Sejnowski, 1995; Fieldtrip implementation Oostenveld et al., 2011) was used to identify and remove components related to blinks and horizontal eye movements. Subsequently to the independent components analysis, epochs that exceeded a signal change of more than 200 µV for any electrode were excluded from analyses.

### Analysis of neural synchronization

In order to investigate whether the periodic repetition of a noise snippet elicited a synchronized response, a fast Fourier transform (FFT) was calculated for each trial that contained a noise-snippet repetition using the data in the 1-s to 3.5-s time window (Hann window tapering; zero-padding). The 1-s to 3.5-s time window was selected because it covered the first 5 snippet repetitions that were present on all trials (cycles 2 to 6; trials without noise repetition were not analyzed due to their low number; N=36, see below). Inter-trial phase coherence (ITPC) was calculated by dividing the complex coefficients that resulted from the FFT by its magnitude and averaging the result across trials (Lachaux et al., 1999). ITPC values in the frequency window ranging from 1.9 to 2.1 Hz were averaged to obtain an ITPC estimate at the 2-Hz stimulation frequency (snippet onset-to-onset interval of 0.5 s). ITPC values were averaged across a fronto-central electrode cluster (F3, Fz, F4, C3, Cz, C4) that is known to be sensitive to neural activity originating from auditory cortex, including neural synchronization (Näätänen and Picton, 1987; Picton et al., 2003; Irsik et al., 2021). To statistically assess whether neural activity was synchronized with the 2-Hz regular rate in the stimulation, a paired samples t-test was used to compare ITPC at 2-Hz to the averaged ITPC across neighboring frequencies (1.8–1.9 Hz and 2.1–2.2 Hz; for a similar procedure see Nozaradan et al., 2011; Nozaradan et al., 2012).

## Experiment 1

Experiment 1 investigates whether neural activity entrained by a noise-snippet repetition self-sustains throughout a brief interruption and the degree to which a noise snippet that reoccurs after the interruption elicits a neural response. Experiment 1 will help understanding how neural representations of frozen noise snippets are maintained.

### Methods

#### Participants

Seventeen participants took part in Experiment 1 (median age: 22 years; range: 18–34 years; 4 male, 13 female).

#### Stimuli and procedure

Participants listened to 4.8-s white-noise sounds in four different conditions. Each condition contained a repetition of a noise snippet, but the number and onset times varied. Conditions are labeled according to the number of snippets embedded in the ongoing noise. Sounds for condition S6 contained six embeddings of a noise snippet, at 0.5, 1, 1.5, 2, 2.5, and 3 s after the onset of the ongoing white noise. For sounds of condition S7, one additional noise snippet was embedded at 3.5 s, and, for sounds of condition S8, two additional noise snippets were embedded at 3.5 and 4 s. Hence, conditions S6, S7, and S8 contained an uninterrupted 2-Hz stimulus rhythm with 6, 7, and 8 cycles, respectively (Figure 2A). Sounds for condition S7* also contained seven embeddings of a noise snippet, but the stimulus rhythm was interrupted and contained one snippet omission in the 7^th^ of the 8 cycles. Snippet onset times for condition S7* were 0.5, 1, 1.5, 2, 2.5, 3, and 4 s (Figure 2A). Comparing activity in the 7^th^ cycle between S6/S7* and S7/S8 allows the investigation of self-sustainability of entrained neural activity. Comparing activity in the 8^th^ cycle between condition S7* and the other three conditions enables the examination of neural sensitivity to a reoccurring noise snippet following a brief discontinuation in the regular, 2-Hz snippet repetition.

**Figure 2:**
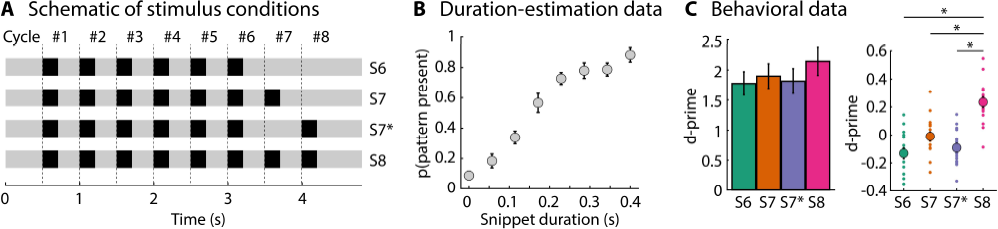
Schematic of stimulus conditions and behavioral data for Experiment 1. A: Schematic of the stimulus conditions. The light gray part represents the longer noise stimulus and black boxes represent the embedded noise snippet. Dashed, vertical lines mark the onsets of the noise snippets for 8 cycles. B: Proportion of ‘pattern-present’ responses as a function of snippet duration in the threshold-estimation procedure. C: Behavioral data from the main experiment. Left: Mean d-prime values across participants. Right: Same data as displayed on the left-hand side, but here the between-participant variance was removed (Masson and Loftus, 2003). Large dots reflect the mean d-prime across participants, whereas the smaller dots reflect the d-prime for each participant. Error bars reflect the standard error of the mean. *p_Holm_ < 0.05

Each participant underwent the procedure to estimate the snippet duration that would yield a rate of 0.75 for the detection of a snippet repetition (described above). For three participants, the targeted detection rate was minimally lowered (but always ≥0.7) because of numerical issues resulting from the logistic function fit used to estimate the threshold. The estimated duration was used throughout the main part of the experiment. Participants listened to six blocks while EEG was recorded. For two participants, EEG data from one block could not be used due to technical issues during recording. Each block comprised 18 trials for each of the four stimulus conditions. A unique, newly created noise snippet was generated for each trial. In addition, 6 trials without any embedded frozen noise snippets were also presented in each block. These ‘pattern-absent’ trials were used to calculate the false alarm rate for each participant. Hence, participants listened overall to 108 trials for each of the four conditions containing a noise-snippet repetition (‘pattern present’) and 36 trials without an embedded noise snippet (‘pattern absent’; for similar procedures see Ng et al., 2012).

#### Analysis of behavioral data

D-prime values were analyzed using a one-way repeated measures analysis of variance (rmANOVA) with the factor Condition (S6, S7, S7*, S8), followed up by planned t-tests and multiple-comparisons correction using Holm’s method. The Inclusion Bayes Factors (BF_incl_) are also provided for the analysis effects based on a Bayesian rmANOVA in JASP software (van den Bergh et al., 2019; JASP, 2022).

#### Neural responses to stimulus cycles

Analyses of neural data focused on stimulation cycle #7 and #8, for which the four conditions differed (all four conditions contained an embedded noise snippet in the first six cycles). Neural activity was averaged across the fronto-central electrode cluster (F3, Fz, F4, C3, Cz, C4) to focus on auditory-cortex responses (Näätänen and Picton, 1987; Picton et al., 2003). Responses to the first five repetitions of a noise snippet (cycles 2–6) show a positive deflection in the 0–0.2-s time window and a negative deflection in the 0.2–0.4-s time window after snippet onset, jointly reflecting the entrained neural activity (Figure 3). To obtain a response estimate separately for cycles #7 and #8, the amplitude in the 0.2–0.4-s time window after cycle onset was subtracted from the amplitude in the 0–0.2-s time window. This resulted in one response-magnitude estimate for each of the four conditions (S6, S7, S7*, S8) and each cycle (#7, #8).

**Figure 3:**
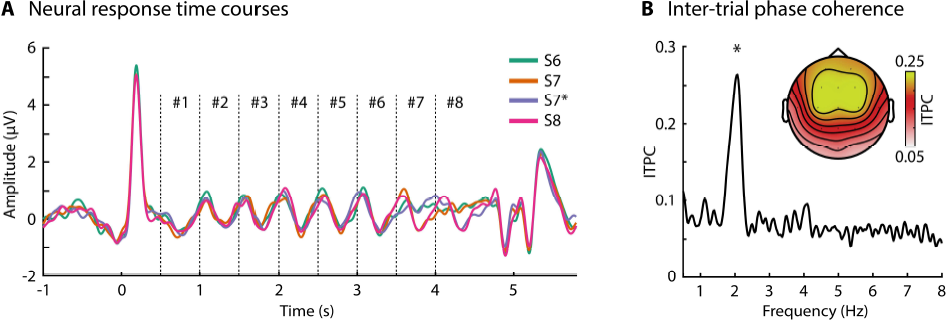
Neural responses to periodic repetitions of embedded noise snippets. A: Neural response time courses (averaged across channels from a fronto-central electrode cluster) for the four conditions time-locked to sound onset. Vertical, dashed lines mark the onsets of the embedded noise snippets. B: Inter-trial phase coherence (ITPC; averaged across channels from a fronto-central electrode cluster) and the topographical distribution of ITPC values for the 2-Hz stimulation frequency. ITPC was calculated across the four conditions containing a noise-snippet repetition (focusing on data from cycles 2-6). *p < 0.05 Analyses for neural activity during cycle #7 tested whether neural activity self-sustains during an interruption of a snippet repetition. The rmANOVA revealed a significant effect of Condition (F_3,48_ = 8.968, p = 8 · 10^−5^, ω^2^ = 0.259, BF_incl_ = 1451). Comparisons show that the response magnitude for S7 and S8 was larger compared to S6 and S7* (S7 vs S6: t_16_ = 3.164, p_Holm_ = 0.024, d = 0.767; S7 vs S7*: t_16_ = 4.983, p_Holm_ = 8.1 · 10^−4^, d = 1.208; S8 vs S6: t_16_ = 3.058, p_Holm_ = 0.024, d = 0.742; S8 vs S7*: t_16_ = 3.578, p_Holm_ = 0.013, d = 0.868). Moreover, the response magnitude was significantly different from zero for S7 (t_16_ = 7.014, p = 2.9 × 10^−6^, d = 1.7) and S8 (t_16_ = 4.038, p = 9.5 × 10^−4^, d = 0.979), but not for S6 (t_16_ = 0.499, p = 0.624, d = 0.121) and S7* (t_16_ = −0.497, p = 0.626, d = 0.121), providing little evidence for self-sustained periodic neural activity despite the hard task that required participants to focus attention on the noise-snippet repetitions that created the stimulus regularity (Figure 4A).

To investigate whether entrained neural activity self-sustains during a brief interruption of a snippet repetition, analyses focused on activity during the 7^th^ cycle (onset at 3.5 s). No noise snippet was presented during the 7^th^ cycle for conditions S6 and S7*, whereas conditions S7 and S8 contained a noise snippet in cycle 7. If entrained activity self-sustains during the interruption of the snippet repetition, a larger than zero response magnitude for the S6/S7* conditions and no response difference between the S6/S7* and the S7/S8 conditions would be expected. If entrained activity does not self-sustain during discontinuation, the S7 and S8 conditions should elicited a larger response than the S6 and S7* conditions, and responses for S6 and S7* may not differ from zero.

To investigate whether a noise snippet reoccurring after the interruption elicits a neural response, analyses focused on neural activity during the 8^th^ cycle (onset at 4 s). Conditions S7* and S8, but not conditions S6 and S7, contained a noise snippet in the 8^th^ cycle. Comparison of responses to S7* relative to S6 and S7 will test whether the auditory system is sensitive to the reoccurrence of the noise snippet, whereas comparison of responses to S7* relative to S8 will test whether the response to the reoccurring noise snippet differs relative to the response to a snippet presented in an uninterrupted sequence.

A one-way rmANOVA with the within-participants factor Condition (S6, S7, S7*, S8) was calculated, separately for cycles #7 and #8, followed up by planned t-tests and multiple-comparisons correction using Holm’s method. The Inclusion Bayes Factors (BF_incl_) are also provided for the rmANOVA effects.

## Results

The rmANOVA for perceptual sensitivity (d-prime) revealed a significant effect of Condition (F_3,48_ = 17.205, p = 1 · 10^−7^, ω^2^ = 0.025, BF_incl_ = 101678). D-prime was larger for S8 compared to S6 (t_16_ = 5.756, p_Holm_ = 1.4 · 10^−4^, d = 1.396), S7 (t_16_ = 4.315, p_Holm_ = 0.002, d = 1.046), and S7* (t_16_ = 6.234, p_Holm_ = 7.2 · 10^−5^, d = 1.512). D-prime was also larger for S7 compared to S6, but only when Holm-correction was omitted (t_16_ = 2.156, p_Holm_ = 0.140, p = 0.047, d = 0.523). S7* did not statistically differ from S6 or S7 (p_Holm_ > 0.24), indicating that there was no benefit from the reoccurring noise snippet for pattern detection.

Inter-trial phase coherence was larger at the 2-Hz snippet-repetition frequency compared to neighboring frequencies (t_16_ = 3.157, p = 0.006, d = 0.766; Figure 3), showing that neural activity synchronized with the periodic presentation of frozen noise snippets. In fact, the amplitude of the neural response to the snippet increased from the first to the second cycle (t_16_ = 3.037, p = 0.008, d = 0.737), but did not further increase from the second to the third cycle (t_16_ = 0.074, p = 0.942, d = 0.018). That only one repetition of the snippet was needed to elicit a full-magnitude response indicates that the brain had some form of memory trace of the running noise prior to the first snippet repetition. The topographical distribution shows largest ITPC values at fronto-central electrodes, suggesting that neural responses originate from auditory cortex (Näätänen and Picton, 1987; Picton et al., 2003).

Analyses for neural activity during cycle #8 tested whether a reoccurring noise snippet following an interruption in the regular repetition elicits a neural response. The rmANOVA revealed a significant effect of Condition (F_3,48_ = 8.433, p = 1.3 · 10^−4^, ω^2^ = 0.240, BF_incl_ = 738). Comparisons focused on the differences in neural response magnitude for the S7* relative to the other conditions. The response magnitude for S7* was greater than the response magnitude for S6 and S7 (S7* vs S6: t_16_ = 2.789, p_Holm_ = 0.039, d = 0.676; S7* vs. 7: t_16_ = 3.260, p_Holm_ = 0.025, d = 0.791), whereas it did not differ from the response magnitude for S8 (t_16_ = 0.686, p_Holm_ = 1, d = 0.166). Moreover, the response magnitude was significantly different from zero for S7* (t_16_ = 3.573, p = 0.003, d = 0.867) and S8 (t_16_ = 5.252, p = 7.9 × 10^−5^, d = 1.274), but not for S6 (t_16_ = −0.257, p = 0.801, d = 0.062) and S7 (t_16_ = −0.75, p = 0.464, d = 0.182). These results demonstrates that the auditory system maintains a memory trace of the specific spectro-temporal structure of a noise snippet embedded in an ongoing noise despite an interruption of a previously established stimulus regularity (Figure 4B).

**Figure 4:**
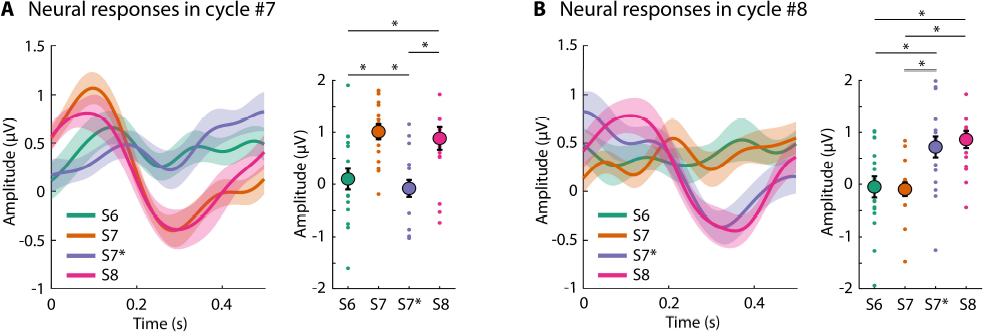
Neural activity in the 7^th^ and 8^th^ stimulation cycle. A: Neural activity in cycle #7. S7 and S8 conditions, but not S6 and S7* conditions, contained a snippet in the 7^th^ cycle. The mean neural response for each condition is depicted in the right plot – the response was calculated as the subtraction of the amplitude in the 0.2–0.4-s time window from the amplitude in the 0–0.2-s time window (left). The large dots reflect the mean amplitude across participants, whereas the smaller dots reflect the mean amplitude for each participant. B: Neural activity in cycle #8. S7* and S8 conditions, but not S6 and S7 conditions, contained a snippet in the 8^th^ cycle. The plots are similar to panel A. Error bars reflect the standard error of the mean. *p_Holm_ < 0.05.

### Summary

The results of Experiment 1 show little evidence that neural activity entrained by a stimulus rhythm self-sustains after the rhythm discontinues (Figure 4A). Critically, a frozen noise snippet reoccurring after a brief interruption elicited a neural response that appeared similar in magnitude compared to the neural response elicited by a noise snippet that was preceded by uninterrupted snippet repetitions (Figure 4B). The results show that, once the auditory cortex establishes a memory trace for the noise snippet, it faithfully responds to another repetition of a noise snippet despite an interruption in rhythmic stimulation.

Stimulation in Experiment 1 was regular at a 2-Hz rate and the snippet reoccurred after the interruption in-sync – that is, at an integer interval – with the preceding stimulus rhythm. A large body of behavioral and electrophysiological work has shown that perceptual and neural sensitivity to a stimulus can depend on its temporal relation to a preceding stimulus rhythm, that is, whether it occurs in-sync or out-of-sync with it (Jones and Boltz, 1989; Large and Jones, 1999; Barnes and Jones, 2000; Jones et al., 2002; Schroeder and Lakatos, 2009; Cravo et al., 2013; Henry and Herrmann, 2014; Henry et al., 2014; Herrmann et al., 2016; Nobre and van Ede, 2018). In Experiment 1, the timing of the reoccurring noise snippet was in-sync with the preceding stimulus rhythm, which perhaps explains the strong response it elicited. Experiment 2 was conducted in order to investigate directly whether the temporal relation to a preceding stimulus rhythm affects neural sensitivity to a reoccurring noise snippet.

## Experiment 2

### Methods and Materials

#### Participants

Twenty individuals participated in Experiment 2 (median age: 21 years; range: 18–34 years; 6 male, 14 female). None of the participants who took part in Experiment 1 took part in Experiment 2.

#### Stimuli and procedure

Participants listened to 5-s white-noise sounds in four different conditions (Figure 5). Each condition contained a repetition of a noise snippet, as in Experiment 1. Sounds for the ‘absent’ condition contained six embeddings of a noise snippet, at 0.5, 1, 1.5, 2, 2.5, and 3 s after the onset of the ongoing white noise (same as S6 in Experiment 1). The label ‘absent’ here refers to the absence of a reoccurring noise snippet after the regular snippet repetition discontinued. The other three conditions contained one additional snippet reoccurring after a brief discontinuation in the stimulus rhythm. In the ‘early’, ‘on-time’, and ‘late’ conditions, the reoccurring snippet was embedded at 3.75, 4, and 4.25 s, respectively. The snippet onset for the ‘early’ and ‘late’ conditions was temporally placed in anti-phase (out-of-sync) relative to the 2-Hz stimulus rhythm established by the preceding 6 snippet embeddings. The snippet onset for the ‘on-time’ condition was placed in-sync/in-phase with the preceding stimulus rhythm (same as 7* in Experiment 1).

**Figure 5:**
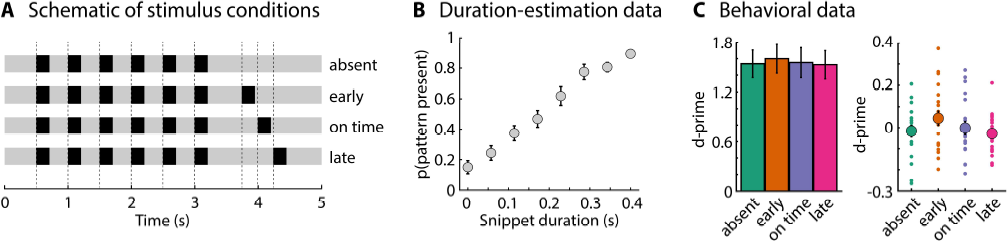
Schematic of stimulus conditions and behavioral data for Experiment 2. A: Schematic of the stimulus conditions. The light gray part represents the longer noise stimulus and black boxes represent the embedded noise snippet. Dashed, vertical lines mark the onsets of the noise snippets. B: Proportion of ‘pattern-present’ responses as a function of snippet duration in the threshold-estimation procedure. C: Behavioral data from the main experiment. Left: Mean d-prime values across participants. Right: Same data as displayed on the left-hand side, but here the between-participant variance was removed (Masson and Loftus, 2003). Large dots reflect the mean d-prime across participants, whereas the smaller dots reflect the d-prime for each participant. Error bars reflect the standard error of the mean. The were no significant differences between conditions.

Similar to Experiment 1, each participant underwent the procedure to estimate the snippet duration that would yield a rate of 0.75 for the detection of a snippet repetition (see descriptions above). For one participant, the targeted detection rate was minimally lowered (but ≥0.7) because of numerical issues resulting from the logistic function fit. The estimated duration was used throughout the main experimental procedures. Participants listened to six blocks while EEG was recorded. Each block comprised 18 trials for each of the four stimulus conditions. As for Experiment 1, a unique, newly created noise snippet was generated for each trial. Six trials without any embedded frozen noise snippets were also presented in each block to calculate the false alarm rate for each participant (Ng et al., 2012). Participants listened overall to 108 trials for each of the four conditions containing a noise-snippet repetition (‘pattern present’) and 36 trials without an embedded noise snippet (‘pattern absent’).

#### Analysis of behavioral data

D-prime values were analyzed using a one-way rmANOVA with the within-participants factor Condition (absent, early, on-time, late). A significant effect was resolved by post hoc comparisons and multiple-comparisons correction using Holm’s method. The Inclusion Bayes Factors (BF_incl_) are also provided for the rmANOVA effects based on a Bayesian rmANOVA in JASP software (van den Bergh et al., 2019; JASP, 2022).

#### Neural responses to sequence-final noise snippet

As for Experiment 1, neural activity was averaged across electrodes of the fronto-central electrode cluster (F3, Fz, F4, C3, Cz, C4), focusing on auditory-cortex responses. An estimate of the response magnitude to the embedded noise snippet was obtained by subtracting the amplitude in the 0.2–0.4-s time window after snippet onset from the amplitude in the 0–0.2-s time window.

Responses were extracted for the ‘early’ (3.75–4.25 s), ‘on-time’ (4–4.5 s), and ‘late’ (4.25–4.75 s) conditions in order to investigate whether the onset-time of the reoccurring noise snippet relative to the preceding stimulus rhythm affects the neural response to the reoccurring snippet. As a control, neural responses were extracted from the same time windows for ‘absent’ trials, during which no snippet reoccurred at 3.75, 4, nor 4.25 s.

A two-way rmANOVA was calculated using the within-participants factors Condition (early, on-time, late) and Snippet Presence (snippet absent, snippet present). A significant Condition effect was resolved by post hoc comparisons and multiple-comparisons correction using Holm’s method. Inclusion Bayes Factors (BF_incl_) are provided for the rmANOVA effects.

## Results

The rmANOVA for the behavioral data was not significant (F_3,57_ = 0.912, p = 0.441, ω^2^ < 0.001, BF_incl_ = 0.174; Figure 5C), that is, behavioral performance did not differ between conditions. The absence of a behavioral effect is consistent with the results from Experiment 1, where a snippet that reoccurred after a brief interruption of a stimulus rhythm also did not lead to a pattern-detection benefit relative to when no snippet reoccurred (S7* vs. S6; Figure 2C).

As for Experiment 1, neural activity synchronized with the regular repetition of embedded, frozen noise snippets. That is, ITPC was larger at the 2-Hz frequency at which snippets were repeated compared to ITPC at neighboring frequencies (t_19_ = 3.325, p = 0.004, d = 0.744; Figure 6). Similar to Experiment 1, the amplitude of the neural response increased from the first to the second snippet (t_19_ = 5.857, p = 1.2 × 10^−5^, d = 1.310), but did not further increase from the second to the third snippet (t_19_ = 0.334, p = 0.742, d = 0.075). The topographical distribution shows largest ITPC values at fronto-central electrodes, again suggesting that synchronized neural activity originates from the auditory cortex (Näätänen and Picton, 1987; Picton et al., 2003). Pooling ITPC data across the two experiments revealed that ITPC at the 2-Hz stimulation frequency (relative to ITPC at neighboring frequencies) was larger for participants who required a longer snippet duration for achieving a similar detection threshold (r = 0.453, p = 0.005; independent of conditions, because the snippet duration was identical across conditions for each participant).

**Figure 6:**
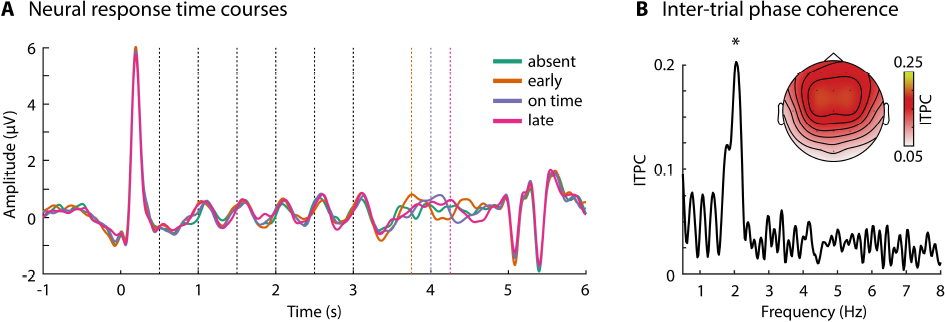
Neural synchronization to periodic repetitions of noise snippets. A: Neural response time courses (averaged across electrodes from a fronto-central cluster) for the four conditions time-locked to sound onset. Vertical, dashed lines mark the onsets of the noise snippets. B: Inter-trial phase coherence (ITPC; averaged across electrodes from a fronto-central cluster) and the topographical distribution of ITPC values for the 2-Hz stimulation frequency. ITPC was calculated across the four conditions containing a noise-snippet repetition (focusing on data from cycles 2-6). *p < 0.05

The noise snippet reoccurring after a brief interruption in the stimulus rhythm elicited a larger neural response than was elicited by the ‘absent’ condition, for which no snippet reoccurred (main effect of Snippet Presence: F_1,19_ = 32.076, p = 1.8 · 10^−5^, ω^2^ = 0.48, BF_incl_ = 606; Figure 7B). There was no effect of Condition (F_2,38_ = 0.488, p = 0.618, ω^2^ < 0.001, BF_incl_ = 0.265) and no Condition × Snippet Presence interaction (F_2,38_ = 2.186, p = 0.126, ω^2^ = 0.04, BF_incl_ = 0.789; Figure 7C, left). In other words, the three conditions containing a reoccurring snippet elicited a response (relative to snippet absence), demonstrating that the auditory system maintains sensitivity to reoccurring noise snippets embedded in a longer noise sound regardless of whether the snippet reoccurs in-sync or out-of-sync with a previous stimulus rhythm. There was also no difference in the response magnitude between the three timing conditions when responses were not normalized relative to the ‘absent’ condition (effect of Condition: F_2,38_ = 1.843, p = 0.172, ω^2^ = 0.019, BF_incl_ = 0.835; Figure 7C, right).

**Figure 7:**
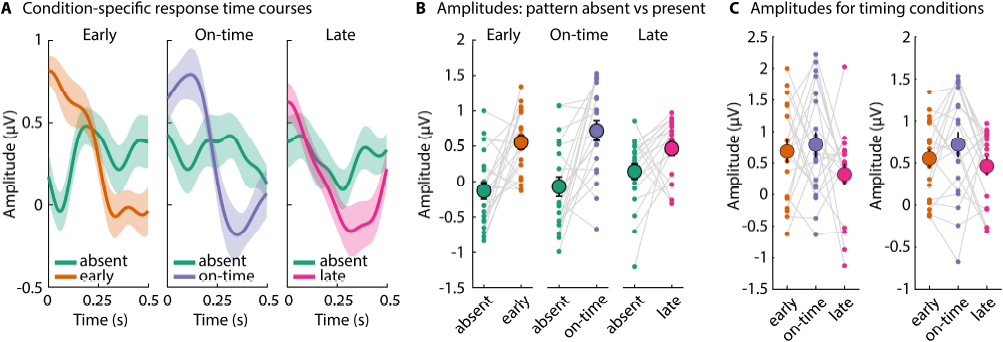
Neural responses to a reoccurring noise snippet at different times following a brief interruption in the regular snippet repetition. A: Response time courses for each condition (early, on-time, late) and the response time course for the ‘absent’ condition for each of the corresponding time windows (early: 3.75–4.25 s; on-time: 4–4.5 s; late: 4.25–4.75 s; x-axis is time-locked to snippet onset for each of these time windows). B: Mean response amplitude, calculated as the subtraction of the amplitude in the 0.2–0.4-s time window from the amplitude in the 0–0.2-s time window. C: Mean response amplitude for the three timing conditions (early, on-time, late) normalized by the ‘absent’ condition (left; the ‘absent’ condition was subtracted from the other three conditions) and non-normalized (right). The large dots reflect the mean amplitude across participants, whereas the smaller dots reflect the mean amplitude for each participant. Error bars reflect the standard error of the mean. Responses were significantly larger for the ‘snippet present’ compared to the ‘snippet absent’ conditions, but there were no significant differences between timing conditions.

## Discussion

The current study investigated how the brain maintains a neural representation of repeating structure in sounds. In two EEG experiments, participants listened to a longer, ongoing white-noise sound which comprised shorter, frozen noise snippets that repeated at a regular rate (Andrillon et al., 2015; Andrillon et al., 2017; Dauer et al., 2022). In several conditions, the snippet repetition discontinued for a brief period after which a noise snippet reoccurred. Results show that neural activity entrained with the regular snippet repetition, but that this periodic neural activity does not sustain during the discontinuation period. Nevertheless, the auditory cortex responds to the reoccurring noise snippet similarly compared to a noise snippet for which the snippet repetition is not discontinued. This response invariance holds regardless of the degree of synchrony of the reoccurring noise snippet relative to the previously established stimulus regularity. Overall, the current study demonstrates that, for the short timescale investigated here, the auditory cortex sensitively responds to a previously learned noise structure in an ongoing sound independent of temporal interruptions.

### Self-sustainability of neural activity

Neural activity becomes entrained by repeating sounds or structure in sounds (Picton et al., 2003; Lakatos et al., 2008; Stefanics et al., 2010; Nozaradan et al., 2011; Thut et al., 2011; Henry and Obleser, 2012; Nozaradan et al., 2012; Lakatos et al., 2013b; ten Oever et al., 2017; Lakatos et al., 2019; Obleser and Kayser, 2019; Herrmann et al., 2023). The peak at 2 Hz in the phase-coherence spectrum of the current study is consistent with entrained activity (Figures 3 and 6). Entrained neural activity – if oscillatory in nature – is hypothesized to self-sustain after periodic stimulation discontinues (Pikovsky et al., 2001; Lakatos et al., 2013b; Henry and Herrmann, 2014; Keitel et al., 2014; Lin et al., 2021; van Bree et al., 2021; Keitel et al., 2022), but there was no evidence for self-sustainability in the current study. That is, there was no clear neural responses in the time window during which the next snippet would have occurred if the repetition had continued (S6 and S7* in Experiment 1; Figure 4). Moreover, neural activity was smaller than activity elicited by a snippet presented in the same time window as part of the snippet repetition (Figures 4; see responses for cycle #7: S6/S7* vs S7/S8; and responses for cycle #8: S6/S7 vs S8).

The current stimulus paradigm may arguably be well suited for the investigation of self-sustained neural activity, assuming its existence, because there are no acoustic edges or changes associated with the occurrence of the noise snippet nor with the discontinuation of the snippet repetition (Figure 1; Agus et al., 2010; Agus and Pressnitzer, 2013; Luo et al., 2013; Andrillon et al., 2015; Rajendran et al., 2016; Andrillon et al., 2017; Dauer et al., 2022). The longer, ongoing noise continues without interruption. Moreover, the detection of the regular repetition of frozen noise snippets is challenging (as indicated by the off-ceiling d-prime values in Figures 2 and 5, and the titration of the detection rate to 0.75) and requires participants to focus attention to the sounds. Attention is likely a facilitator for entrained neural activity and self-sustainability (Lakatos et al., 2013b; Lakatos et al., 2013a; Henry and Herrmann, 2014; Ringer et al., 2023a). Since the current study finds little evidence for self-sustainability, this adds to the ongoing discussion about whether entrained neural activity self-sustains when a stimulus periodicity discontinues (Lakatos et al., 2013b; Lin et al., 2021; van Bree et al., 2021). The absence of self-sustained activity may further raise the question of whether neural activity entrained by the stimulus regularity reflects neural oscillatory activity or event-evoked activity (Klimesch et al., 2007; Sauseng et al., 2007; Capilla et al., 2011; Fellinger et al., 2011; Notbohm et al., 2016; Keitel et al., 2021). Self-sustained activity would not be expected for event-evoked activity. The response to a snippet reoccurring after the regular repetition had discontinued was very similar to the response to the snippet when it was part of the regular repetition (S7* vs S8 in Experiment 1), which is in line with event-evoked neural activity in the current study.

### Invariant neural response to the reoccurring frozen noise snippet

The current study shows that the neural response to a frozen noise snippet, reoccurring after a regular snippet repetition had briefly discontinued, is robust and invariant to a brief discontinuation (Figure 4). That is, the response to the reoccurring noise snippet did not differ from the response to a noise snippet for which the regular repetition was not discontinued previously. The response was further invariant to changes in the temporal relation between the onset of the reoccurring snippet relative to the previous stimulus regularity (Figure 7). Overall, it appears that once a neural representation for a specific noise snippet is established, the brain responds to the reoccurrence of the snippet invariantly, at least over the short timescale investigated here.

Several previous works have shown that individuals better detect a noise snippet repetition when the snippet that is repeated recurs across trials compared to a snippet that does not recur (Agus et al., 2010; Agus and Pressnitzer, 2013; Andrillon et al., 2015; Andrillon et al., 2017; Dauer et al., 2022). Previous work further shows that individuals show behavioral correlates of memory for random noise or tone structure in sounds even days and weeks after exposure (Agus et al., 2010; Viswanathan et al., 2016; Bianco et al., 2020; Bianco et al., 2023). That the brain responds with similar magnitude to a reoccurring noise snippet following an interruption compared to a snippet that is part of a regular, non-interrupted sequence is consistent with a memory representation for a noise snippet that lasts long once the representation has been established.

### Relation between neural responses and behavioral detection performance

The results of the current study show that participants are more sensitive to the presence of a repetition of a frozen-noise snippet in an ongoing noise when the number of regular, uninterrupted repetitions increases (Figure 2C; Experiment 1). However, an additional reoccurring noise snippet following a brief interruption in the repetition did not improve detection performance (Figures 2C and 5C). The absence of a benefit for repetition-detection was observed for reoccurring snippets that were in- and out-of-sync with the preceding stimulus regularity. While some previous works have shown perceptual benefits for in-sync compared to out-of-sync stimuli (Jones et al., 2002; Jones et al., 2006; Hickok et al., 2015; Herrmann et al., 2016), other recent research has failed to find such a perceptual benefit (Bauer et al., 2015; de Graaf and Duecker, 2022; Lin et al., 2022; Sun et al., 2022). Stimulus designs often differed substantially among previous studies, making it hard to pinpoint under which conditions perceptual benefits for in-sync compared to out-of-sync stimuli can be observed.

Although the brain responded robustly to the reoccurring noise snippet, this apparently did not translate to a behavioral performance increase (Figures 2C and 5C). Moreover, neural responses to the frozen noise snippet were already present for the first snippet repetition (cycle #2) and did not further increase with the number of repetitions, whereas behavioral detection performance increased with the number of repetitions (Figure 2C). Other work also suggests that one repetition of a noise snippet is sufficient for the brain to elicit a strong response, although this was not strictly analyzed previously (Andrillon et al., 2015; Ringer et al., 2023a). Moreover, neural entrainment to periodic sound stimulation has been observed even for sub-threshold sound stimulation (e.g., tone beeps in noise at a low signal-to-noise ratio; ten Oever et al., 2017). This mismatch between the pattern of neural responses and the behavioral patterns perhaps suggests that a memory trace of the noise snippet is established relatively quickly and stably, whereas the detection of the repetition requires another, possibly higher cognitive process involving decision making.

## Conclusions

The current study investigated how the brain maintains representations of repeating structure in sounds. Participants listened to a longer, ongoing white noise sound which comprised shorter, frozen noise snippets that repeated at a regular rate. The current study aimed to answer whether neural activity self-sustains during a brief interruption of the repetition and how the brain responds to a noise snippet that reoccurs after the interruption. No evidence for self-sustained neural activity during the interruption was observed. However, the auditory cortex responded to a reoccurring noise snippet with similar magnitude compared to a noise snippet for which the snippet repetition had not been interrupted. This response invariance was observed for different onset times of the reoccurring noise snippet relative to the previously established regularity. The current study thus demonstrates that the auditory cortex sensitively responds to a previously learned noise structure in an ongoing sound independent of temporal interruptions.

## Acknowledgements

We thank Christie Tsagopoulos for her help with data collection for both experiments. The research was supported by the Canada Research Chair program (CRC-2019-00156, 232733) and the Natural Sciences and Engineering Research Council of Canada (Discovery Grant: RGPIN-2021-02602).

